# Human adipose-derived mesenchymal stromal cells from face and abdomen undergo replicative senescence and loss of genetic integrity after long-term culture

**DOI:** 10.1101/2021.04.16.440228

**Authors:** Priscilla Barros Delben, Helena Debiazi Zomer, Camila Acordi da Silva, Rogério Schutzler Gomes, Fernanda Rosene Melo, Patricia Dillenburg-Pilla, Andrea Gonçalves Trentin

**Author notes:** Corresponding Author: Andréa Gonçalves Trentin, Department of Cell Biology, Embryology, and Genetics, Federal University of Santa Catarina, Brazil, Campus Universitário, 88040-900, Trindade, Florianópolis, SC, Brazil. Priscilla Barros Delben and Helena Debiazi Zomer have contributed equally to this work. Authors information: Priscilla Barros Delben Helena Debiazi Zomer Camila Acordi da Silva Rogério Schutzler Gomes Fernanda Rosene Melo Patricia Dillenburg-Pilla.

## Abstract

Body fat depots are heterogeneous concerning their embryonic origin, structure, exposure to environmental stressors, and availability. Thus, investigating adipose-derived mesenchymal stromal cells (ASCs) from different sources is essential to standardization for future therapies. *In vitro* amplification is also critical because it may predispose cell senescence and mutations, reducing regenerative properties and safety. Here, we evaluated long-term culture of human facial ASCs (fASCs) and abdominal ASCs (aASCs) and showed that both met the criteria for MSCs characterization, but presented differences in their immunophenotypic profile, and differentiation and clonogenic potentials. The abdominal tissue yielded more ASCs, but facial cells displayed fewer mitotic errors at higher passages. However, both cell types reduced clonal efficiency over time and entered replicative senescence around P12, as evaluated by progressive morphological alterations, reduced proliferative capacity, and SA-β-galactosidase expression. Loss of genetic integrity was detected by a higher proportion of cells showing nuclear alterations and γ-H2AX expression. Our findings indicate that the source of ASCs can substantially influence their phenotype and therefore should be carefully considered in future cell therapies, avoiding, however, long-term culture to ensure genetic stability.

## INTRODUCTION

Mesenchymal stromal cells (MSCs) are promising candidates for application in cell therapy and regenerative medicine due to their broad paracrine effects on tissue-resident cells combined with their low immunogenicity when allogenic transplanted [1]. MSCs reside in virtually all adult tissues, playing important roles in homeostasis and repair [2]. To characterize this heterogeneous cell population, the International Society for Cellular Therapy (ISCT) specified guidelines for defining MSCs that include plastic adherence, fibroblastic morphology, mesodermal differentiation potential, mesenchymal expression (CD73, CD90, and CD105), and absence of hematopoietic (CD34 and CD45) markers [3,4]. However, despite sharing the minimum stem cell requirements, MSCs from different niches were described to display distinct characteristics *in vitro* and in pre-clinical studies *in vivo* [5–8].

Our research group has been studying the biology and therapeutic potential of human MSCs from different tissues and body sources, including periodontal ligament, placenta, facial and abdominal dermis, and adipose tissue [7–11]. We previously compared the procedures for MSC isolation from abdominal adipose tissue and abdominal dermis and found that while the first source has a greater cellular yield, the latter has improved ability to promote the closure of skin wounds *in vitro* and in pre-clinical mouse models [7,8]. These studies confirmed that MSCs derived from different tissues present distinct capabilities and highlighted the need to characterize the different sources of MSCs in various pre-clinical and clinical contexts to find the most appropriate source for each application.

Among the various tissue sources of MSCs, the adipose tissue stands out by its great availability and accessibility in addition to higher cellular yield, proliferative rate, and genetic stability *in vitro* when compared to other sources [7,12–15]. However, adipose tissues are not equal, and although most body depots originate from the mesoderm, the facial adipose tissue partially originates from the ectodermal neural crest [16–19]. Further, while subcutaneous adipose abdominal deposits are composed of two layers of tissue, one dense (superficial and vascularized) and the other lamellar (deeper and less vascularized), the facial adipose tissue is irregular and composed of only one tissue layer. [17,20]. Moreover, the facial adipose tissue is highly exposed to harmful environmental stressors, such as temperature, ultraviolet radiation, pollution, and synthetic products [21], and these factorsmay influence the biology of their MSCs (adipose-derived MSCs, hereafter called ASCs). Moreover, while ASCs from the abdominal adipose tissue (aASCs) are obtained by elective liposuctions or abdominoplasties and are widely characterized, facial ASCs (fASCs) can be obtained by elective rhytidectomies, an obscure source of ASCs [22–24].

Because millions of cells are required for cellular therapy transplantation [25,26], the selection of the ideal source of MSC needs to consider not only the characteristics of their stem cell but also their availability for clinical use. A large number of MSCs can be achieved through extensive *in vitro* amplification, which may predispose the accumulation of cellular damage associated with senescence, and reduce its regenerative properties and therapeutic safety [27–30]. Therefore, ensuring the genetic integrity of MSCs during long-term culture is essential to establish safe protocols for therapeutic applications.

A previous study by Kim and co-workers (2013) [31] revealed that MSCs of abdominal and facial (eyelid) adipose tissues exhibit a distinct pattern of cell surface antigen expression, as well as different potentials of proliferation, differentiation, and telomerase activity. However, comparative information regarding the effects of long-term culture on facial and abdominal adipose-derived MSCs remains elusive. To this end, we compared MSCs isolated from these depots regarding the replicative senescence and genetic integrity after prolonged *in vitro* expansion, which can influence the efficiency and safety of these cells for future cellular therapies in regenerative medicine.

## RESULTS

### Characterization

Human subcutaneous ASCs were isolated from facial and abdominal fat depots. To confirm that the isolated cells could be classified as MSCs following the ISCT guidelines [3], flow cytometry analysis was performed and revealed the expression of hematopoietic markers CD34 and CD45 in less than 1.3% of cells. In contrast, the mesenchymal markers CD73, CD90, and CD105 were found in more than 81.7% of both ASCs (Figure 1). Interestingly, significantly greater expression of CD90 (98.3±0.3%, p=0.0013) and CD105 (94.3±0.6%, p=0.04) was found in fASCs compared with aASCs (93±0.5% and 81.7±4.1%, respectively).

**Figure 1:**
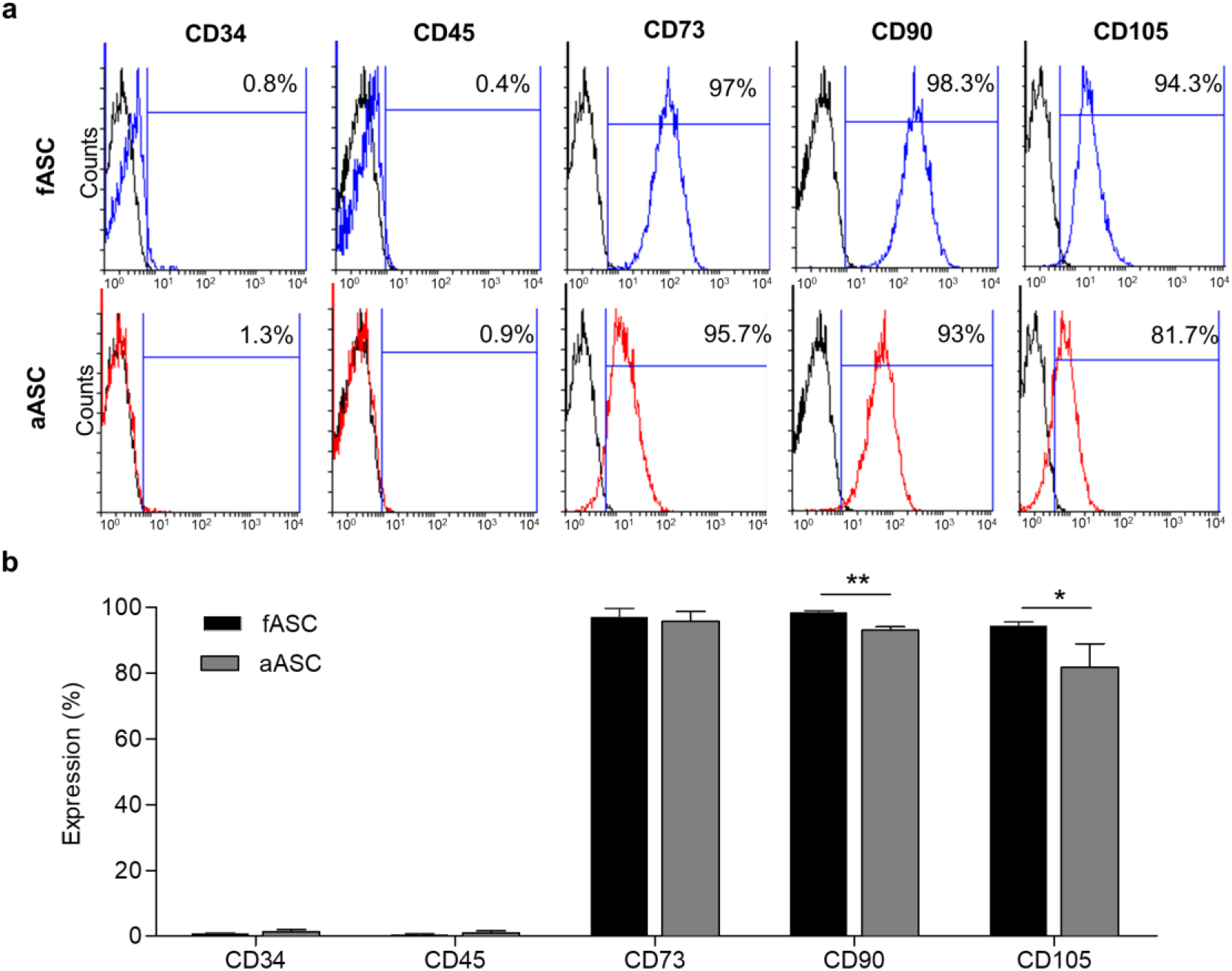
ASCs from face and abdomen express CD73, CD90 and CD105 and do not express CD34 and CD45. a) Representative histograms of flow cytometry to CD34, CD45, CD73, CD90, and CD105. b) Increased expression of CD90 and CD105 in fASCs compared with aASCs. * p<0.05 and ** p<0.01 by unpaired t-test.

Next, we comparatively evaluated the mesodermal differentiation potential of ASCs, and both were able to differentiate into adipocytes, osteocytes, and chondrocytes when cultured in specific inductive media (Figure 2). Adipocyte-like cells, identified by lipid inclusions stained by Oil Red Sudan lipid 5B (Figure 2a, b), appeared 3 days earlier (day 6) in aASC cultures than in fASCs (day 9). aASCs presented a sustained increased adipogenic differentiation potential during the 21 days of culture compared to fASCs (2.4-fold at 21 days, p<0.0001, Figure 2b). On the other hand, the osteogenic differentiation, assessed by the presence of calcified extracellular matrix, stained by Alizarin Red (Figure 2c), and quantified by the intensity of the color (Figure 2d), was initially observed at day 21 in fASC culture, 3 days earlier than in aASCs (at day 24). Nevertheless, both ASCs presented similar levels of osteogenic differentiation at day 27 (the endpoint of this study) (Figure 2d). Finally, we comparatively evaluated the chondrogenic differentiation potential of ASCs, and both formed similar cartilaginous-like pellets after 28 days of culture in the specific inductive media (Figure 2e, f). However, increased deposition of collagen fibers (based on the intensity of Mallory staining) was detected in fASCs compared to aASCs (1.58-fold, p<0.0001).

**Figure 2:**
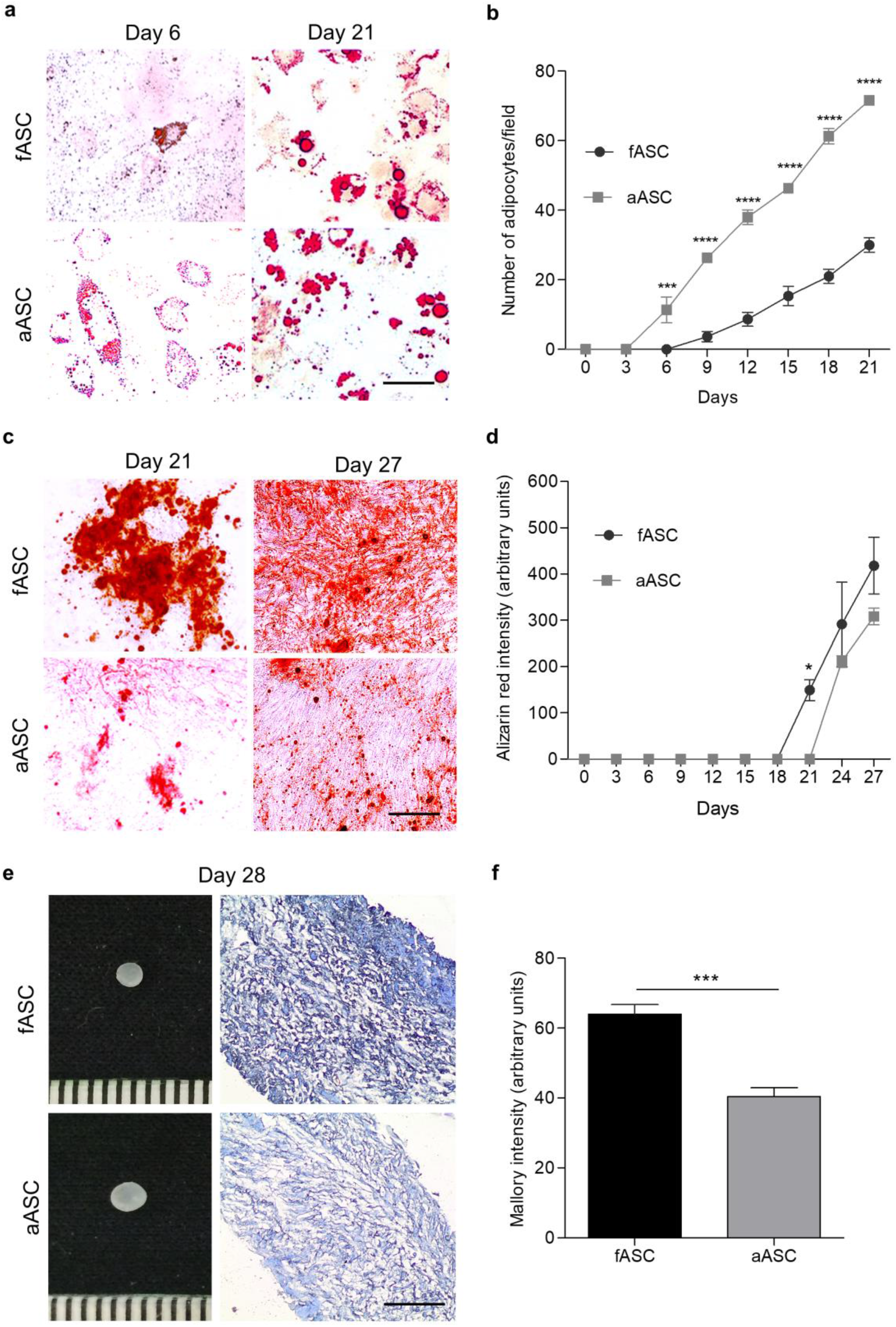
Abdominal ASCs show increased adipogenesis while facial ASCs display higher osteogenic and chondrogenic potential. a) Adipogenic differentiation attested by lipid droplets stained by Oil Red Sudan lipid 5B at days 6 (left panel) and 21 (right panel) of culture in the specific differentiation medium. (b) Quantification of stained cells (adipocytes) during the 21 days of culture. (c) Osteogenic differentiation visualized by extracellular calcium matrix stained by Alizarin Red at days 21 (left panel) and 27 (right panel) of culture in specific differentiation medium. d) Quantification of the staining intensity during the 27 days of culture. (e) fASCs and aASCs formed 2-mm cartilaginous-like pellets after 28 days in the chondrogenic differentiation medium (left). Pellets were histologically processed, and the cartilaginous extracellular matrix (collagen) was detected with Mallory staining and (f) the intensity quantified at day 28. For (b), (d), and (f), values represent the mean ± SEM of three biological replicates (three donors) in four fields/replicate. *** p<0.001 and **** p<0.0001 by (b, d) two-way ANOVA or (f) unpaired t-test. Scale Bar 200 μm.

Despite the differences found between fASCs and aASCs concerning the CD90 and CD105 expression and the differentiation performance, their immunophenotypic profile and mesodermal differentiation potential demonstrate that both exhibit the characteristics of MSCs established by ISCT. Therefore, these findings confirm that ASCs can be harvested from both abdominal and facial adipose tissue depots, but display some distinct characteristics *in vitro*, which may potentially influence their applicability *in vivo*.

### Cellular yield, growth curve, and clonogenicity

We next comparatively examined the applicability of the face and the abdomen as potential sources of MSCs to future clinical assays by evaluating the cellular yield upon isolation. Strikingly, we were able to harvest 29-fold more cells per gram of tissue (40 ± 2.28×10^4^) from the abdomen than from the face (1.4 ± 0.08 ×10^4^) (Figure 3a). Because a greater proliferation rate *in vitro* may overcome an initial lower cellular yield, we next investigated the proliferative potential of these cells and found that they share similar growth curves and population doubling times at P2 (Figure 3b, c). These findings indicate that larger numbers of ASCs can be obtained from the abdominal adipose tissue compared to the facial depot, suggesting a clinical availability advantage of aASCs.

**Figure 3:**
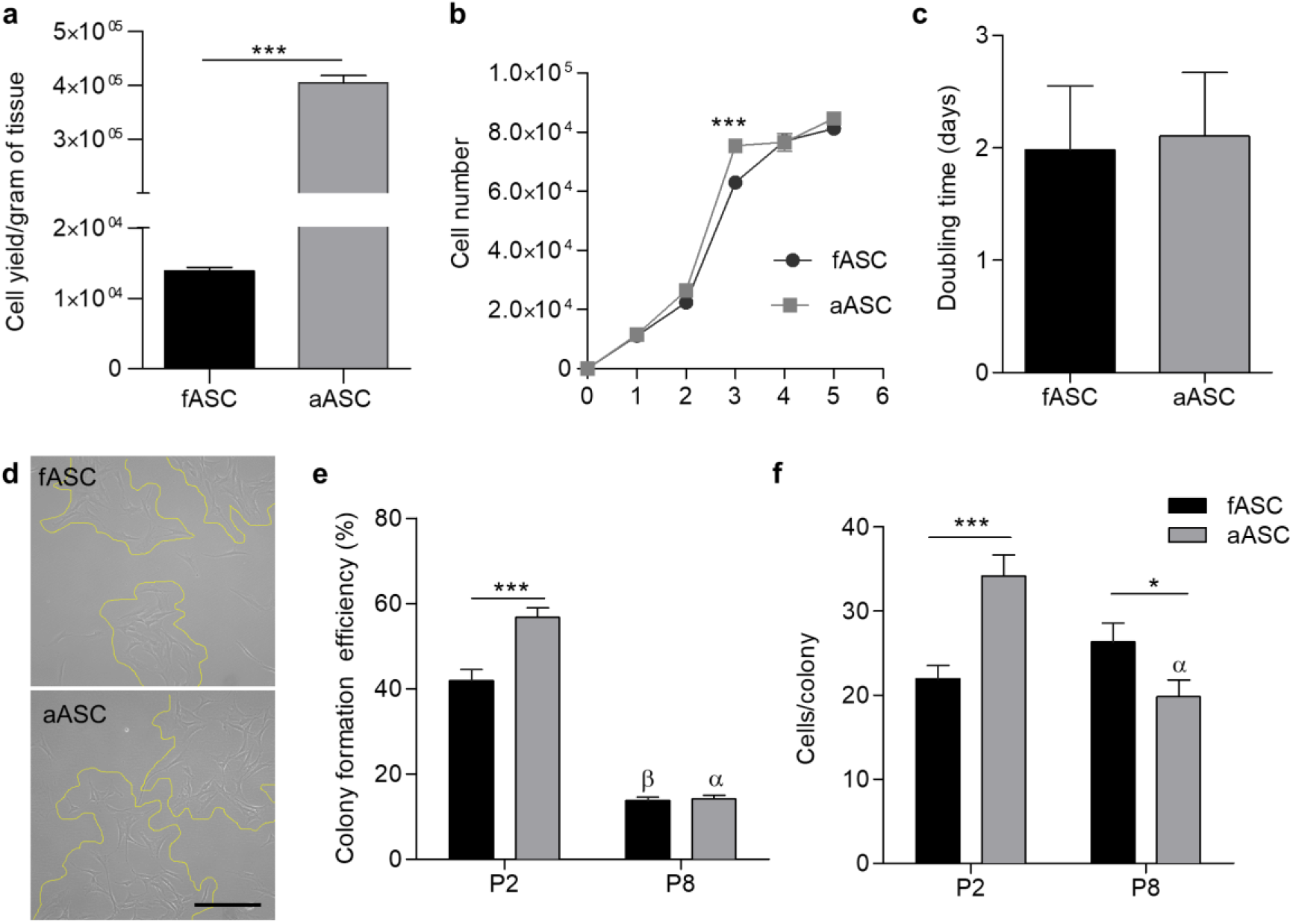
Cellular yield and proliferative profile of ASCs. (a) Cellular yield per gram of tissue (adherent cells after 24h of culture). (b) Growth curve at P2. (c) Population doubling time at P2. (d) Representative photomicrographs of colony-forming units at P2. Yellow lines highlight colony boundaries. Scale Bar 200μm). (e) Colony-forming unit efficiency and (f) number of cells per colony at P2 and P8. For all graphs, bars represent the mean ± SEM of three donors. * p<0.05, ** p<0.01, *** p<0.001 by unpaired t-test (fASC x aASC). α and β indicate statistical significance between different passages of the indicated cell type by unpaired t-test.

We then evaluated the *in vitro* clonogenicity of ASCs (Figure 3d-f). The colony-forming unit assay assumes that only stem/progenitor cells can form colonies, and therefore, the clonal efficiency is indicative of their proportion whereas the colony size (number of cells per colony) suggests the proliferative potential of the founder stem/progenitor and/or its daughter cells in culture [32]. Increased cloning efficiency (1.3-fold) and colony size (1.5-fold) at P2 were found in aASC (56.7±7.8% and 34.1±9.3 cells/colony) compared to fASC cultures (41.8±9.5% and 21.9±5.8 cells/colony) (Figure 3e, f). These findings indicate that through the isolation and culture conditions used here, aASC cultures have more cells with stem/progenitor phenotypes which are more proliferative than fASCs at low passages. Moreover, the clonal efficiency was greatly reduced to below 20% at P8 in both ASCs (14.1 ± 3% in aASCs and 13.8 ± 2.7% in fASCs, a 4-fold and 3-fold reduction compared to P2, respectively), suggesting a loss of this stem cell characteristic over time in culture. The colony size was also reduced at P8 in aASCs (1.7-fold compared to P2), whereas in fASCs, similar values were observed between these passages.

Together, these findings show that a higher amount of cells can be harvested from the abdominal fat by the isolation procedures used here than from the facial depot. Although a greater proportion of aASCs exhibit the stem/progenitor cell phenotype in P2, fASCs and aASCs share similar proliferative profiles at lower passages and have their clonogenicity reduced after a few passages *in vitro*. Notably, fASCs kept their proliferation rate stable over the passages (P2-P8), which raised the interesting hypothesis that fASCs might be more resistant to replicative senescence *in vitro*.

### Evaluation of cellular growth, senescence, and genetic integrity over time in culture

It is widely accepted that massive amplification of MSCs ultimately results in replicative senescence *in vitro* [33]. Therefore, we next investigated if fASCs and aASCs were differentially affected by replicative senescence after prolonged *in vitro* expansion (Figure 4).

**Figure 4:**
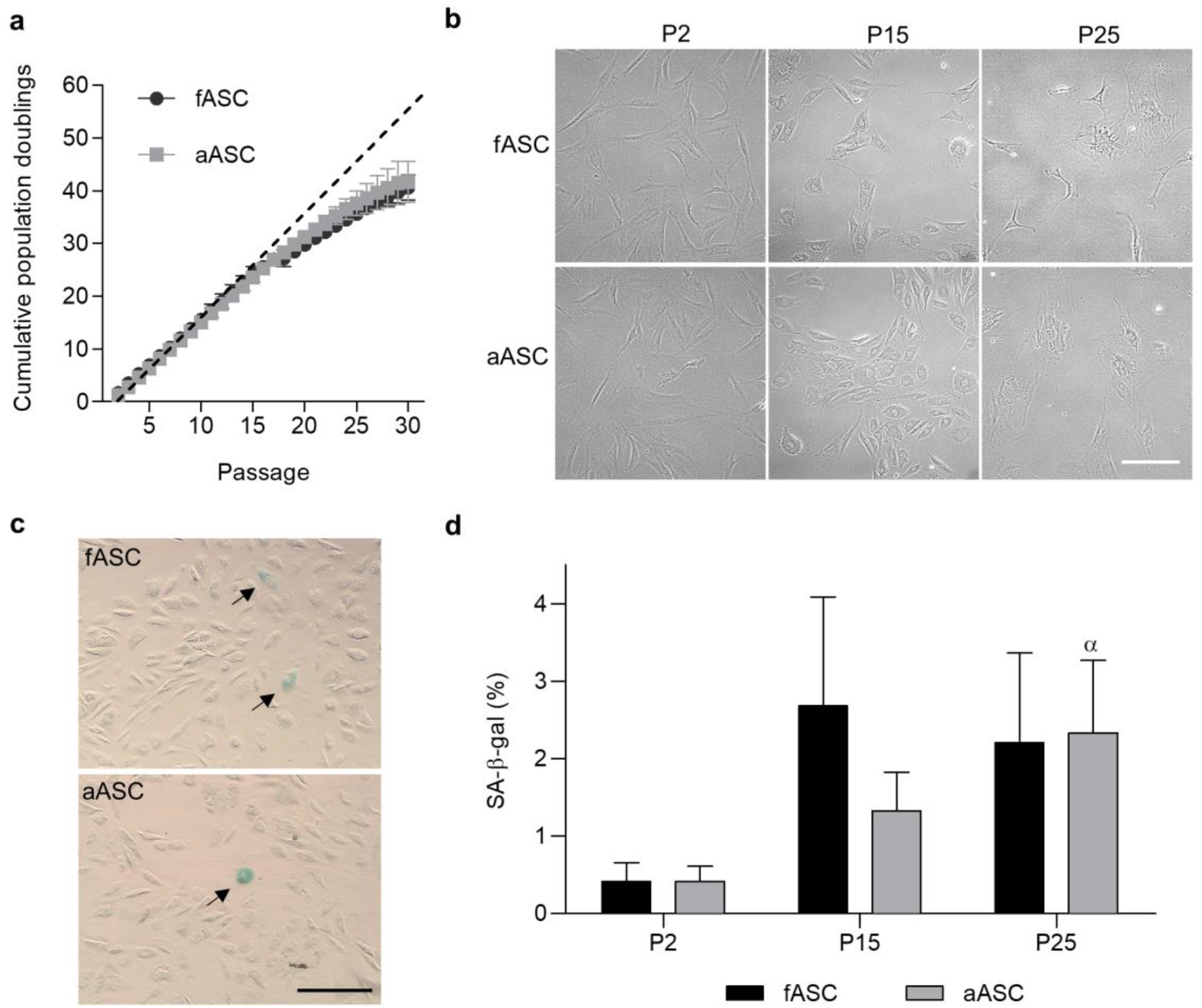
Long-term culture of ASCs leads to replicative senescence. (a) Cumulative population doublings during 30 passages. (b) Representative phase-contrast photomicrographs of fASCs and aASCs at low (P2), medium (P15), and high (P25) passages. (c) Representative phase-contrast photomicrographs of senescence-associated β-galactosidase (SA-β-gal) staining (arrows) at low (P2), medium (P15), and high (P25) passages. Scale Bar 200μm. (c) Quantification of SA-β-gal-positive cells. Bars represent the mean of four different fields containing a minimum of 200 cells in 3 biological replicates (3 donors) ± SEM. α indicates statistical significance between P2 and P25 for aASCs.

Evaluation of the cumulative population doubling showed that the proliferative capacity of both ASCs was similarly reduced after P15, corresponding to 25 population doublings (Figure 4a). Morphological alterations were observed in both cell types, progressively changing from a spindle shape at low passages (P2), to a hypertrophic, enlarged, flattened, and irregular morphology with cytoplasmic granules and extensions (characteristics of senescent cells) at medium (P15) and high (P15) passages (Figure 4b), suggesting that replicative senescence takes place at similar time points in both ASCs.

To further evaluate cell senescence, we quantified cells positively stained for senescence associated-beta-galactosidase (SA-β−gal), a biological marker of cellular senescence and aging [34] (Figure 4c, d). Interestingly, less than 5% of SA-β−gal-positive ASCs were recorded during all evaluated passages (P2-P25), despite the increased proportion from lower (0.4 ± 0.8% and 0.4 ± 0.6% of fASCs and aASCs, respectively), to medium, (2.6 ± 4.6% and 1.3 ± 1.7% of fASCs and aASCs, respectively) and high passages (2.2 ± 4% and 2.3 ± 3.2% of fASCs and aASCs, respectively). Although no differences were observed between cell types, considerable variation was noted among donors, which resulted in a large standard deviation. Significant values across passages were recorded only for aASCs (P2 *vs*. P25, p=0.038); however, a trend was observed for fASCs (P2 *vs*. P15, p=0.592), suggesting that both ASCs gradually undergo replicative senescence over time in culture.

Replicative senescence is directly associated with the accumulation of DNA damage secondary to mitosis [35]. Mitotic errors can disrupt genomic integrity, leading to cellular transformation, which impairs therapeutic effects and raises safety issues [15]. Therefore, to evaluate possible cellular damages secondary to mitosis in our ASC cultures, the frequency of nuclear alterations was assessed by cytokinesis-block micronucleus assay (Figure 5a, b). Micronuclei, an indication of breakage and/or loss of chromosomes [36], was not observed in either cell type or any evaluated passage (P5-P25). On the other hand, nuclear buds, a marker of gene amplification and/or elimination of the DNA repair complexes [36], were found in a similar small, relative frequency at low passage (P5) in both cell cultures (0.14 ± 0.06 for fASCs and 0.09 ± 0.08 for aASCs) (Figure 5a). At high passage (P25), however, the frequency of nuclear buds increased in both ASCs compared to P5 (0.27 ± 0.06 for fASCs, p=0.001, and 0.20 ± 0.12 for aASCs, p=0.04). Besides, small, relative frequencies of nucleoplasmic bridges, a nuclear alteration associated with DNA strand break misrepair and/or telomere end-fusions [36], were also similar in the ASC cultures at P5 (0.14 ± 0.17 for fASCs and 0.26 ± 0.13 for aASCs), values 2.2-fold (p=0.02) and 2-fold (p=0.0001), respectively, increased at P25. At this high passage, significantly greater frequency of nucleoplasmic bridges was found in aASCs (0.53 ± 0.08) than fASCs (0.32 ± 0.14) (Figure 5b). Together, these findings indicated that nuclear alterations occurred in low proportion at initial passages with gradual increase over time in culture in both ASCs. Notably, fASCs displayed fewer errors associated with DNA strand break misrepair and/or telomere end-fusions than aASCs.

**Figure 5:**
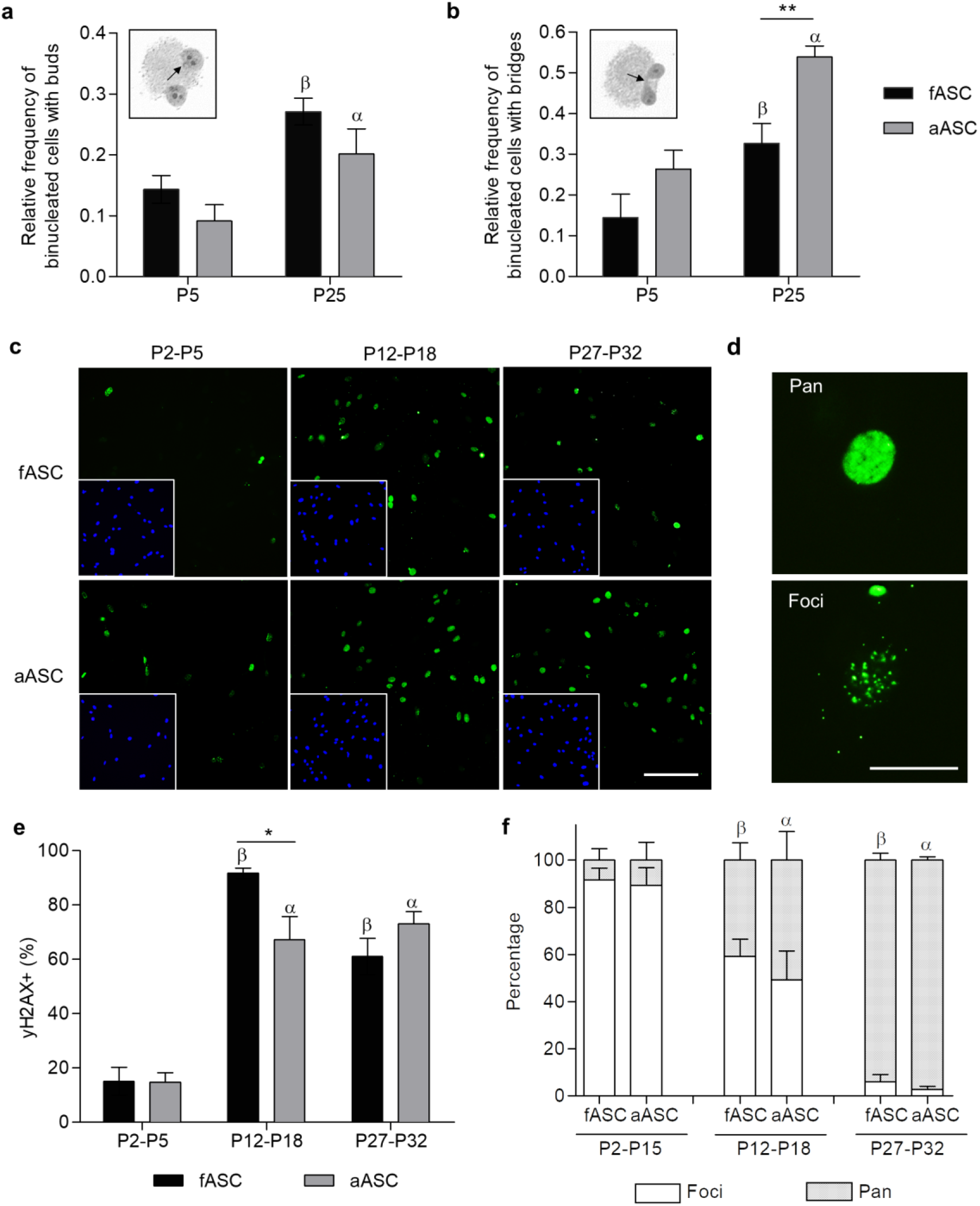
Long-term culture induces loss of genetic integrity in ASCs. Relative frequency of binucleated cells with (a) buds and (b) bridges at low (P5) and high (P25) passages. Representative photomicrographs of the corresponding nuclear abnormalities are shown in graph insets. Bars represent the mean of 3 biological replicates ± SEM. β and α indicate statistical significance between P5 and P25 fASCs and aASCs, respectively. ** p<0.01 by unpaired t-test. (c) Representative immunofluorescence photomicrographs of fASCs (upper panel) and aASCs (lower panel) at low (P2-P5), medium (P12-P18), and high (P27-P32) passages. Cell nuclei were counterstained with DAPI (insets). Scale Bar 200 μm. (d) Representative pictures of the γ-H2AX pan (top panel) and focal (lower panel) expression. Scale Bar 25 μm. (e) Quantification of γ-H2AX positive cells. Bars represent the mean of 3 biological replicates ± SEM. * p<0.05 by unpaired Student t-test (fASCs x aASCs at P12-P18). α and β indicate statistical significance among different passages for the indicated cell type. (f) Percentage of cells displaying pan and focal nuclear patterns of γ-H2AX expression. No difference was observed between cell types by unpaired Student t-test. α and β indicate statistical significance (p<0.05 by unpaired Student t-test) among different passages for the indicated cell type.

Next, the genetic integrity of ASCs during prolonged culture was evaluated by the expression of γ-H2AX, a sensitive indirect marker of DNA double-strand breaks secondary to genetic damage [37]. Immunofluorescence staining revealed 15.1 ± 14.6% of fASCs and 14.8 ± 10.4% of aASCs positives to γ-H2AX at low passages (P2-P5), with no difference between them (Figure 5c-e). These values were significantly increased at medium (P12-P18) (91.6 ± 5.5% and 67.2 ± 25.7% of fASCs and aASCs, respectively) and high (P27-P32) passages (61.1 ± 20% and 73 ± 13.5% of fASCs and aASCs, respectively). Significantly greater expression of γ-H2AX was found in fASCs than aASCs (p=0.019) at P12-P18, despite their similar values at P27-P32. Moreover, two patterns of nuclear γ-H_2_AX staining were observed: foci- and pan-expression (Figure 5d). The increased number of foci ultimately leads to pan-expression of γ-H2AX and directly correlates to the degree of DNA damage [37]. Interestingly, both ASCs presented mostly the foci pattern of γ-H2AX expression at low passages, with a progressive shift to the pan-expression throughout the medium and high passages (figure 5f), suggesting a similarly progressive activation of cellular responses to DNA damage over time in culture.

Taken together, these data indicate that the prolonged culture leads to replicative senescence and loss of genetic integrity in both ASCs. Noteworthy, fASCs seemed to be more resistant to these effects considering their stable colony size between P2 and P8 and their lower frequency of nuclear aberrations, namely nucleoplasmic bridges.

## DISCUSSION

In this study, we comparatively characterized the mesenchymal phenotype and the effects of long-term culture on ASCs isolated from human facial and abdominal subcutaneous adipose tissue. We hypothesized that the ASCs derived from these tissues would display distinct characteristics *in vitro*. We showed that both fASCs and aASCs met the minimum criteria established by the ISCT for mesenchymal stem cell characterization [3], but differed in their immunophenotypic profile, differentiation, and clonogenic potentials (Figure 6). Nevertheless, both ASCs underwent replicative senescence and loss of genetic integrity after long-term culture. Our findings indicate that the source of ASCs can substantially influence their phenotype and therefore should be carefully considered in future cell therapies, avoiding, however, long-term culture to ensure genetic stability.

**Figure 6:**
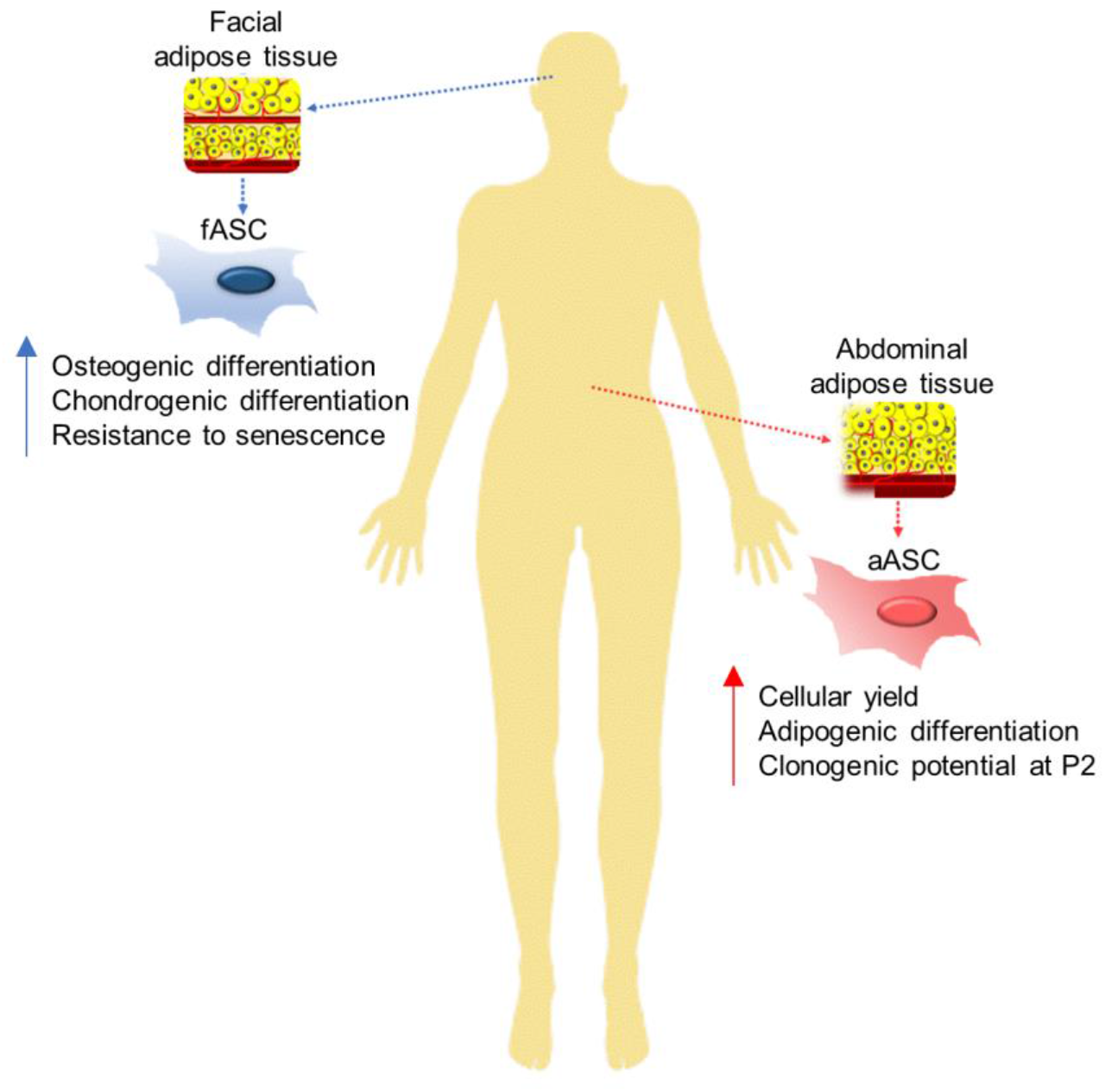
Human ASCs from face and abdomen have different properties. ASCs derived from the facial subcutaneous adipose tissue depot have greater chondrogenic and osteogenic potential than abdominal ASCs. Besides, long-term culture seems to affect the fASCs less than their mesodermal counterparts. On the other hand, the abdominal adipose tissue is more abundant, and cell isolation procedures yields a greater amount of aASCs, which in turn have improved adipogenic potential and greater clonogenicity in early passages. These findings indicate that the source of ASCs can substantially influence their phenotype and therefore should be carefully considered in future cell therapies.

Considering the wide variety of tissues available as sources of MSCs, their proper characterization is essential for clinical-grade standardization [38]. Here, we showed that despite both ASCs express mesenchymal markers are absent of hematopoietic markers, fASCs present greater expression of CD90 and CD105 compared to aASCs. These findings contrast with those of Kim and co-workers (2013), in which facial (eyelid) ASCs showed a markedly lower expression of CD90 than abdominal ASCs. The functional consequences of different levels of expression of mesenchymal markers remain to be elucidated.

Both ASCs were able to generate osteocytes, chondrocytes, and adipocytes *in vitro*, as previously described by Kim et al. (2013). Through our study of differentiation by time point, we show that fASCs have a higher capacity for osteogenic and chondrogenic differentiation, while abdominal cells have better adipogenic potential *in vitro*. Similarly, studies have demonstrated that other progenitors derived from the neural crest, such as MSCs of the dental pulp and eyelid adipose tissue, have higher chondrogenic and osteogenic potentials when compared with their mesoderm derived counterparts [31,39–41]. These findings were functionally confirmed *in vivo* by a higher efficiency in bone repair, suggesting a promising application of neural crest-derived MSCs in regenerative medicine approaches for bone and cartilage injuries [39,40].

The reduced adipogenic potential of fASCs compared to aASCs observed could be related to the fact that while the facial adipose tissue is dramatically reduced with aging, the abdominal subcutaneous adipose tissue remains an essential long-term energy storage depot in normal adults [17,42]. Morever, ASCs isolated from old individuals have reduced adipogenic potential compared to ASCs from young subjects [43]. In this context, it is noteworthy that although the tissue donors of our study were all adults (32-61 years old), those of facial adipose tissue were on average, 21 years older than the abdominal fat subjects. Therefore, the age difference between fASCs and aASCs could potentially play a role in our findings.

Because cellular therapies require the transplantation of a great number of cells [25,26], the cellular yield and the *in vitro* long-term proliferation and genetic stability are critical characteristics when selecting sources of MSCs. Here, we show that more cells can be harvested per gram of abdominal adipose tissue than the facial counterpart. Considering that the abdominal fat availability is also markedly higher, this deposit has an undeniable advantage when considering the starting cellular yield. Interestingly, no differences were observed in the growth curve and in the doubling time between fASCs and aASCs in P2, suggesting that they display a similar proliferative rate at initial passages. However, different patterns in their ability to form colonies, a characteristic of stem cell self-renewal capacity [32], were observed. Although the aASCs showed higher clonal efficiency and larger colony size than the fASCs in P2, both showed a marked decrease in clonogenicity in P8, indicating a reduction in the stem cell phenotype after a few *in vitro* passages. However, in contrast to the aASCs, the fASC colonies’ size did not change from P2 to P8, suggesting higher consistency in the proliferation rate over passages. This is particularly interesting because the chronological age of the donor has been reported to compromise the potential for self-renewal and proliferation of adult stem cells during *in vitro* expansion [27,44,45]. Thus, given the older age of facial adipose tissue donors in our study, it was expected that the proliferative profile and self-renewal capacity of fASCs would be decreased in comparison to aASCs. These findings indicate that the long-term culture can distinctly affect ASCs.

Continued *in vitro* expansion of MSCs might predispose the accumulation of senescence-associated cellular damage, reducing the regenerative properties of stem cells and their therapeutic safety [27–30]. However, the replicative senescence secondary to massive amplification *ex vivo* cannot be avoided due to the need for a large number of cells for pre-clinical and clinical applications [26]. Therefore, we cultured ASCs for up to 32 passages to comparatively investigate their senescence and genetic integrity. Interestingly, despite different embryonic origins, exposure to stressors, and donor age, fASCs and aASCs were considerably similar after long-term culture. Replicative senescence, evaluated by the presence of hypertrophic and flat cells, reduction of population doublings, and increased activity of SA-β-gal, started around P15 for both cell types.

To investigate if the replicative senescence of ASCs was associated with cellular damage, nuclear aberrations secondary to mitotic errors were identified by cytokinesis-block micronucleus assay [36]. Results revealed a small proportion of nuclear alterations at lower passages (P5) similar between fASCs and aASCs. However, a significant increase in the percentage of nucleoplasmic bridges, a sign of DNA strand break misrepair, and/or telomere end-fusions [36], and in the nuclear buds, a marker of gene amplification and/or elimination of DNA repair complexes [36], was detected in both long-term ASC cultures, indicating they accumulate genetic errors secondary to massive mitosis over time. Noteworthy, nuclear aberrations were less frequent in fASCs, suggesting that they are less affected by prolonged *in vitro* expansion. This finding corroborates with the observation that fASC sustained number of cells per colony over passages, further indicating they may be more stable in culture.

To ensure tissue homeostasis *in vivo*, MSCs conserve their long-term proliferation potential [33]. Therefore, adult stem cells have an exceptionally high ability to repair DNA damages associated with replication and those induced by exogenous genotoxic factors [33]. The first response of cells to DNA breaks is the formation of γ-H2AX DNA repair foci, indicating proper damage recognition [33]. In our study, both ASCs shifted from similarly low levels of foci γ-H2AX expression at initial passages to most of the cells pan-expressing it at P12 and progressively thereafter. The γ-H2AX shift from foci to pan expression over passages is a strong indicator of response pathways’ activation to persistent DNA damage that ultimately induces cell senescence [46,47]. Interestingly, despite the disparity in cell donors chronological age, no major differences were observed between their genetic stability *in vitro*, as initially expected.

In our study, under standard culture conditions, both ASCs had their stem cell phenotype reduced after P5-P8 and showed signs of replicative senescence around P12 when widespread molecular damage response pathways were activated. Because culture conditions can significantly influence the biological properties of MSCs [15], improved culture methods need to be developed to ensure their safety after extensive *in vitro* amplification. As an example of that, Li et al. (2017) expanded ASCs in a serum-free medium according to good manufacturing practices guidelines, and found that although cell proliferation decreased significantly after 15 passages, no evident chromosomal aberrations or relevant gene expression alterations were observed until passage 20, suggesting the protocol is safe and suitable for clinical translation [43,48]. Moreover, another study found that hypoxic conditions improve the DNA repair responses of long-term cultured bone marrow-derived MSCs, but not from ASCs, indicating that culture protocols need to be optimized for each MSC, depending on their tissue origins [25].

Therefore, the ideal culture conditions to ensure the long-term genetic stability of ASCs remain to be determined.

This is the first study comparing human ASCs derived from adipose tissue depots of different embryonic origins, neural crest (ectoderm) and mesoderm, focusing on their genetic integrity after long-term culture. Although previous studies have compared ASCs from different fat depots, most of them focused on mesodermal-derived tissues, such as visceral versus subcutaneous adipose depots [4,49,50]. Furthermore, few studies addressed the safety of cultured human ASCs by evaluating their genomic stability [43]. Nevertheless, our study is not free of limitations. Because the facial and abdominal adipose tissue samples were obtained from different donors, we cannot exclude the possibility of individual variations. The different average ages of the donors may introduce yet another variable. It is also important to consider that the face is more subjected to the burden of biological aging due to its exposure to environmental stressors, such as solar radiation, the main cause of photoaging [24,27,28,51]. However, the subcutaneous adipose tissue is naturally protected by several layers of epidermal cells and dermal tissue, so the influence of environmental stressors might not be functionally relevant on fASCs [52]. Last, the harvesting procedure of adipose tissue can affect cellular viability, yield, proliferation, and stemness. For example, it has been described that ASCs isolated from fat obtained by liposuction present higher proliferation and later senescence than those obtained by surgical resection [43,53]. Therefore, besides their distinct embryonic origins, other factors might have influenced the results observed in our study and should be considered.

In conclusion, our study showed that ASCs derived from the facial subcutaneous adipose tissue depot (originated from the neural crest – ectoderm) have higher CD90 and CD105 expression and greater chondrogenic and osteogenic potential than the abdominal ASCs (originated from the mesoderm). Additionally, long-term culture seems to affect the fASCs less than their mesodermal counterparts, although this finding needs to be further investigated in future studies. On the other hand, abdominal adipose tissue is more abundant, and cell isolation procedures result in a greater amount of aASCs, which in turn have greater clonogenicity in early passages. However, both ASCs show signs of replicative senescence and loss of genetic integrity over time in culture. These findings shed light on the distinct abilities of MSCs derived from different sources and highlight the neural crest-derived ASCs as promising candidates for future bone and cartilage regeneration studies. Ideally, long-term culture under standard methods should be kept at a minimum (up to P5) to ensure genetic integrity. Because *ex vivo* expansion of MSCs cannot be avoided, further studies should investigate culture methods to maintain their phenotypic and functional characteristics, and genomic stability to ensure the safety of future cell therapies in regenerative medicine.

## MATERIAL AND METHODS

### Donors

Human subcutaneous adipose tissue was obtained from healthy (with no chronic, viral, metabolic, ischemic, autoimmune, or other diseases) female donors undergoing elective rhytidectomy (facelift, ages 57, 59, and 61 years old, average 59±2 years old) or abdominal liposuctions (ages 32, 36 and 45 years old, average 38±6 years old) at Dr. Carlos Correa Hospital Plastic Surgery Center (Florianopolis, Brazil). To avoid major individual donor variations, only adults (32-61 years old) with skin type II or III (Fitzpatrick scale) and a body mass index of 18.5-29.9 kg/m^2^ (normal to overweight) were included in this study. Patients consented to participate under the guidelines established for human clinical trials by the human research ethics committee from the Federal University of Santa Catarina (protocol 131.512).

### Isolation and culture of ASCs

Isolation and culture of ASCs were performed as previously described, with few modifications [7]. Briefly, 2 g of subcutaneous facial or abdominal adipose tissue were obtained, sectioned, and incubated with 1% collagenase I (Sigma-Aldrich, USA) for 40 min at 37°C. The enzymatic digestion was blocked by the addition of 10% fetal bovine serum (FBS, Vitrocel, Brazil), and the obtained suspension was filtered through a 70-μm mesh strainer (BD Bioscience) and centrifuged at 300 g for 5 min. The pellet was suspended in erythrocyte lysis buffer (155 mM NH_4_Cl, 12 mM NaHCO_3_, 0.1 mM EDTA), incubated for 10 min at 37°C, and centrifuged again. The supernatant was discarded, and the pellet was suspended in a standard medium consisting of Dulbecco’s Modified Eagle’s Medium (DMEM, Sigma-Aldrich, USA) supplemented with 10% FBS, streptomycin (10 μg/ml; Sigma-Aldrich), and penicillin (200 U/ml; Sigma-Aldrich). Cells were seeded in 25 cm^2^ culture flasks (Corning, USA) and maintained at 37°C in humidified 5% CO_2_. After 24 h, the culture was washed with DMEM, and the medium was changed. This procedure was repeated daily during the first 3 days of culture and every 3 days thereafter.

ASCs in the primary culture (passage zero, P0) were harvested by trypsinization (0.05% trypsin/0.02% EDTA) at 90% confluence. Cells were replated in the standard medium at the initial density of 4×10^3^ cells/cm^2^. The procedure was repeated when monolayers reached 90% of confluence for the next passage (P). ASCs in several passages (P2 to P32) were cryopreserved in 10% dimethyl sulfoxide (DMSO) and 90% FBS at a density of 1×10^6^ cells/ml per cryotube (Sarstedt, Germany) and maintained in liquid nitrogen for further analysis.

### Flow cytometry

Flow cytometry was performed to detect the antigenic markers CD34, CD45, CD73, CD90, and CD105 as previously described [7,54]. Briefly, 1×10^5^ ASCs at P3 were harvested by trypsinization and incubated for 1 h in the dark, at 4°C, with the following fluorochrome-conjugated antibodies: anti-CD34 (PE/clone 581), anti-CD45 (FITC/clone HI30), anti-CD73 (PE/clone AD2), anti-CD90 (FITC/clone 5E10) and anti-CD105 (PercP/clone 266, all antibodies were from BD Bioscience). Fractions of white blood cells were used as positive controls for CD34 and CD45, and IgG isotype antibodies conjugated to fluorophores as negative controls. Cells were washed with phosphate-buffered saline (PBS) and assessed by flow cytometry (FACScalibur; BD Biosciences). Data were analyzed by Flowing Software (Turku Centre for Biotechnology, University of Turku, Finland).

### Adipogenic differentiation

Adipogenic differentiation was performed as previously described, with few modifications [7,9]. Briefly, 90% confluent ASC monolayers at P3-P5 were grown in the standard medium supplemented with 10^−2^ M dexamethasone, 100 μM indomethacin, 2.5 μg/ml insulin, and 0.5 mM isobutyl methylxanthine (all from Sigma Aldrich). Cells were maintained at 37°C and 5% CO_2_ for up to 21 days, with medium changed every 3 days. Cells were fixed with 4% paraformaldehyde and stained with a 2% Oil Red Sudan lipid 5B (Sigma Aldrich) for 10 min at different time points to quantify differentiated cells containing red-colored lipid vacuoles.

### Osteogenic differentiation

Osteogenic differentiation was also performed as previously described, with few modifications [7,9]. Briefly, ASCs at P3-P5 were seeded at a density of 10^4^ cells/well in 24-well plates and maintained in the standard medium until 100% confluence and then transferred to osteogenic inductive medium consisting of standard medium supplemented with 10^−8^ M dexamethasone, 5 μg/ml ascorbic acid, and 3.15 mg/mL β-glycerophosphate (all from Sigma Aldrich). Cells were maintained at 37°C and 5% CO_2_ for up to 28 days, with medium changed every three days. Cells were fixed with 4% paraformaldehyde and stained with 2% Alizarin Red solution (Sigma Aldrich) for 10 min at different time points to observe orange-red colored calcium deposits. The staining intensity was quantified using ImageJ (National Institutes of Health) after conversion to the grayscale and normalization to the total culture dish area.

### Chondrogenic differentiation

For chondrogenic differentiation, 2.5×10^5^ of ASCs at P3-P5 were cultured as a pellet in a 15 mL conical tube in a specific medium (StemXVivo Human / Mouse Kit, R & D Systems) at 37°C and 5% CO_2_. After 28 days, the chondrogenic aggregates were photographed and then fixed (4% paraformaldehyde), histologically processed, sectioned in 5 µm cross-sections, and stained with Mallory (1% Phosphotungstinic acid, 2% Orange G and 0.5% Aniline blue) for 30 min. The intensity of Mallory staining was quantified using ImageJ as previously described [8].

### Cellular yield

To comparatively investigate the initial cellular yield of ASCs obtained from facial versus the abdominal adipose tissue, cells were isolated and cultured for 24 h as described above. Then, floating cells were washed out with PBS, and attached cells were detached by trypsinization and quantified using a hemocytometer.

### Growth curve and doubling time

To compare the fASC and aASC proliferative profiles, 1×10^4^ cells in P2 were plated in DMEM containing 10% BFS in 24-well plates. The number of cells per well was counted daily until cells reached 100% confluency at day 5. The population doubling time was calculated using the formula x = [log10(NH/NI)]/log10(2)], where NI is the inoculum cell number (10^4^) and NH the number of cells harvested.

### Colony Forming Units Assay

The colony-forming units assay was performed as previously described, with minor modifications [7,55]. Briefly, ASCs at P2 and P8 were seeded at low density (50 cells/cm^2^) and cultured for 5 days. Cells were then fixed in 4% paraformaldehyde and stained for 10 min with 2% crystal violet. The number of cells per colony (for colonies with 10 or more cells) and the clonal efficiency (proportion of plated cells that formed colonies) were counted.

### Long-term culture of fASCs and aASCs

To investigate the effects of long-term culture in fASCs and aASCs, cells were cultured for up to 30 passages. Cell morphology was evaluated in all passages by phase-contrast microscopy at 100x, 200x, 400x magnifications (Olympus DP71), and images were taken with a capture system (Olympus MVX10).

### Cumulative population doubling time over passages

The population doubling time was calculated as previously described with minor modifications [56,57]. Briefly, 5×10^4^ cells were seeded per well of a 6-well culture plate and cultured for 3 days, and then they were detached, counted, and re-plated at the same initial density. The procedure was repeated until the number of recovered cells was lower than that of the initial plating. The number of cells per well counted at each endpoint was applied to the doubling time formula. The number of population doublings in each passage was added to the previous endpoint’s number to yield the cumulated doublings.

### Senescence-associated beta-galactosidase activity

Cells at low (P2), medium (P15), and high (P25) passages were fixed with 4% paraformaldehyde for 40 min, washed 3 times with PBS, and stained with Senescence Cells Histochemical Staining kit (Sigma-Aldrich) for 6 h at 37°C in a humidified atmosphere at 5% CO_2_, and processed according to the manufacturer’s instructions. Positive stained cells were evaluated by phase-contrast microscopy at 200x magnification (Olympus DP71), and images were taken with a capture system (Olympus MVX10). The percentage of stained cells was counted in 4 different fields (minimum of 200 cells per biological replicate).

### Cytokinesis-Block Micronucleus assay

Nuclear stability (DNA damage) over time in culture was evaluated by the cytokinesis-block micronucleus assay, as described previously [36,58]. Briefly, 70% confluent ASCs monolayers at P5 or P25 were incubated with 5 mg/ml cytochalasin B (Sigma Aldrich) for 48 h at 37°C and 5% CO_2_. Cells were detached, centrifuged, and the cell pellets were treated for 3 min with a hypotonic solution (0.075% potassium chloride and 1% of DMEM in distilled water). After, cells were fixed for 30 min (90% methanol and 10% acetic acid solution), mounted on glass slides, stained with 0.5% Giemsa (7 min), and analyzed under a light microscope. Six hundred binucleated cells of each replicate were scored for the relative frequency of micronuclei, nucleoplasmic bridges, and nuclear buds.

### γ-H2AX Immunocytochemistry

Double strand DNA breaks were evaluated by immunostaining to γ-H2AX, a marker of DNA damage, as previously described [59]. Briefly, ASCs at low (P2-P5), medium (P12-P18), and high (P27-P32) passages were cultivated under standard conditions for 3 days. Cells were fixed with 4% paraformaldehyde for 30 min at room temperature, permeabilized in PBS containing 0.25% Triton X 100, and unspecified sites were blocked with 10% FBS in PBS. Samples were then incubated with a primary antibody anti-phospho-histone H2AX (γ-H2AX, 1:200, Millipore, USA) overnight at 4°C, followed by incubation with the secondary antibody anti-rat Alexa Fluor 488 (1:500) for 60 min at room temperature. Cell nuclei were stained with DAPI (1:1000, 4’
s-6-diamidino-2-phenylindole, Sigma Aldrich, USA) for 10 min at room temperature. Cells were visualized using an inverted fluorescence microscope (Olympus, DP71), and images were captured at 100x and 400x magnification (Olympus DP83 camera). Cells displaying at least one γ-H2AX focus and irregular labeling of distinct points in the nucleus were considered positives and quantified.

### Statistical Analyses

Statistical analyses were performed using GraphPad Prism software (GraphPad Software, La Jolla, CA, USA). All experiments were performed with 3 biological replicates (3 donors) per ASC type. Gaussian distribution was confirmed by the Shapiro-Wilk test. Then, data were analyzed by unpaired Student’s t-test or two-way analysis of variance (ANOVA) followed by Bonferroni post-hoc test for multiple comparisons, when appropriate. Differences between mean values were considered significant when p < 0.05.

## ACKNOWLEDGEMENTS

We thank all patients who agreed to donate their disposable surplus liposuction and rhytidectomy to this work. We thank nurses Silvana Soares Francisco and Marcelo Conte for mediating our contact with patients. We thank Dra. Débora Cornélio for cytokinesis-block assay training. We thank Dr. Emily Brehm for revising the language of this manuscript. We thank the Multi-User Laboratory for Studies in Biology (LAMEB) of the Federal University of Santa Catarina for the technical support. This work was supported by the Ministry of Science, Technology, Innovations, and Communications / National Development Council Scientific and Technological (MCTIC / CNPq / Brazil), Nacional Institute of Science and Technology in Regenerative Medicine (MCTIC / CNPq/ INCT-REGENERA), grant numbers 456928/2013-8, 465656/2014-5, 407734/2018-8 and 305202/2019-7, Coordination for the Improvement of Level Personnel Superior (CAPES, Brazil) finance code 001. PBD, HDZ, ACS, and FRM received fellowships from CAPES and PDP from CNPq.

